# Reverse-breaking CFS (rev-bCFS): Disentangling conscious and unconscious effects by measuring suppression and dominance times during continuous flash suppression

**DOI:** 10.1101/2024.08.21.608242

**Authors:** Tommaso Ciorli, Lorenzo Pia, Timo Stein

## Abstract

Breaking continuous flash suppression (bCFS) is a widely used experimental paradigm that exploits detection tasks to measure the time an initially invisible stimulus requires to escape interocular suppression and access awareness. One pretty contentious and unresolved issue is whether differences in detection times reflect unconscious or conscious processing. To answer this question, here we introduce a novel approach (i.e., reverse-bCFS [rev-bCFS]) that measures the time an initially visible stimulus requires to be suppressed from awareness. Results from two experiments using face stimuli indicate that rev-bCFS can capture conscious effects, which indicates that contrasting standard bCFS with rev-bCFS can isolate unconscious processing occurring specifically during bCFS. For example, while face inversion impacted both bCFS and rev-bCFS, effects were larger in bCFS, suggesting a distinct contribution of unconscious processing to the advantage of upright over inverted faces in accessing awareness. Combining standard bCFS and rev-bCFS may offer a fruitful approach able to disentangle conscious and unconscious effects occurring during interocular suppression.

## Introduction

When our eyes are exposed to different images, conscious perception does not unify the two images in a unique percept but, rather, it dynamically alternates the two images. Such binocular rivalry (hereinafter BR) is useful for investigating which factors determine competition for visual awareness. Typically, stimuli that dominate perception for a longer time are believed to be consciously prioritized by the visual system (Alpers & Pauli, 2006), whereas those escaping suppression faster to gain dominance are thought to be unconsciously prioritized (Jiang et al., 2007). Over the past two decades, continuous flash suppression (CFS; (Tsuchiya & Koch, 2005)), a variant of BR, has been widely used to investigate visual processing outside awareness, that is in the suppression phase (Pournaghdali & Schwartz, 2020; Sterzer et al., 2014). In CFS, one eye is exposed to a high-contrast dynamic mask (typically updating at 10 Hz) which can suppress a stimulus shown to the other eye for prolonged periods, without the occurrence of perceptual alternations that characterize standard BR.

The so-called “breaking CFS” (bCFS; for reviews, see (Gayet et al., 2014; Stein, Hebart, et al., 2011; Stein, 2019)) paradigm uses simple detection tasks to measure the time an initially suppressed stimulus needs to overcome suppression and access awareness. The rationale is that CFS specifically disrupts conscious processing but allows information to be processed unconsciously. Thus, stimuli that are detected faster are thought to enjoy unconscious prioritization, boosting the sensory signal into consciousness. Differences in detection times in bCFS have been taken as evidence that a wide range of higher-level cognitive processes can occur unconsciously (for a review, see (Hassin, 2013)): facial and bodily features (Ciorli & Pia, 2023; Lanfranco et al., 2023), semantic content (Costello et al., 2009; Jiang et al., 2007; Kerr et al., 2017; Yang & Yeh, 2011), emotions (Hedger et al., 2015; Vetter et al., 2019; Yang et al., 2007; Zhan, Hortensius, et al., 2015), degree of familiarity (Geng et al., 2012; Gobbini et al., 2013; Stein et al., 2014, 2016), threat (Gayet et al., 2016), multisensory information (Aller et al., 2015; W. Zhou et al., 2010), food (Ciorli et al., 2024; Lee et al., 2022), and abstract concepts (Sklar et al., 2012). These results have challenged the conventional view according to which high-level processing during BR had been thought to be restricted to dominant phases only while having little influence on suppression times (Blake & Logothetis, 2002).

However, some have advocated caution on the validity of bCFS to reveal unconscious processing (Hesselmann & Moors, 2015; Lanfranco et al., 2021; Moors et al., 2017; Stein & Sterzer, 2014). Indeed, bCFS relies on responses to a subjectively *visible* stimulus, an approach in stark contrast with classic dissociation techniques in which unconscious processing is demonstrated when an *invisible* stimulus continues to influence behavior (Kouider & Dehaene, 2007; Schmidt & Vorberg, 2006). In bCFS, detection differences do not necessarily reflect unconscious processing under CFS but they could, alternatively, reflect differences in conscious stimulus processing during the transition into awareness, including differences in decision criteria. One common approach to exclude such conscious effects is to contrast bCFS with a non-CFS control condition, where the same stimuli are superimposed on the masks, thus not inducing interocular suppression. Most studies did not find bCFS-like detection effects in non-CFS control conditions (e.g., (Costello et al., 2009; Jiang et al., 2007; Mudrik et al., 2011; W. Zhou et al., 2010); for a review, see (Stein, 2019)), which has been taken as evidence that bCFS effects must have reflected unconscious processing under CFS.

However, it has become increasingly clear that non-CFS control conditions are not suitable to control for conscious effects. First, empirical results indicate that they are not sensitive enough to pick up effects that many other psychophysical procedures reliably reveal, such as the face inversion effect. Second, non-CFS control conditions are perceptually vastly different from the CFS-like interocular dynamics characterized by perceptual uncertainty, and unpredictability of target appearance. With greater uncertainty, differences in decision criteria could have a larger effect on differences in detection times, and thus spuriously amplify bCFS effects. As mimicking CFS-induced perceptual uncertainty without interocular suppression is extremely difficult to achieve, one solution would be a control condition that controls for conscious effects but also involves interocular suppression.

### The reverse-breaking CFS proposal

Here, we introduce “reverse-breaking continuous flash suppression” (hereinafter, rev-bCFS), which reverses the standard bCFS trial sequence. Specifically, at the beginning of a trial the target stimulus is consciously perceived before it is gradually suppressed by the mask until the mask fully suppresses the target. Participants press a key as soon the last cue of the stimulus disappears from awareness (contrasting with standard bCFS where participants detect target appearance). In other words, rev-bCFS measures the time it takes for a stimulus to be suppressed (i.e., the transition from conscious to unconscious, or conscious disappearance) – that is, how long it persists and dominates conscious perception, as compared to bCFS that measures the time it takes for a stimulus to overcome suppression (i.e., the transition from unconscious to conscious). As they involve similar interocular suppression, bCFS and rev-bCFS are better comparable than bCFS and standard non-CFS control conditions. The key difference lies in the type of information processing occurring before the response, that is, unconscious processing before the subjective visibility threshold in bCFS, and interocular conscious processing preceding the same threshold in rev-bCFS. The comparison of bCFS vs. rev-bCFS could thus allow to measure and control for the contribution of conscious effects to bCFS detection differences.

We tested this approach in two experiments using face stimuli. In Experiment 1, we compared the well-established face inversion effect (FIE) between bCFS and rev-bCFS. The FIE reflects the visual system’s enhanced sensitivity to process faces with an upright (i.e., in their prototypical spatial representation) compared to an inverted (i.e., rotated 180°) orientation. In addition to bCFS, better detection of upright faces has been observed in a wide variety of tasks, including visual search, attentional blink, and backward masking (Garrido et al., 2008; Lewis & Edmonds, 2003; Lewis & Edmonds, 2005; Rossion et al., 1999; Tyler & Chen, 2006; Van Belle et al., 2015), but, curiously, typically not in the classic non-CFS control condition (e.g., (Jiang et al., 2007), but see (Stein, Hebart, et al., 2011)). We considered the FIE as a candidate for an effect that may involve conscious and unconscious mechanisms (Stein & Peelen, 2021). A larger effect in bCFS than in rev-bCFS would thus provide tentative evidence for unconscious face processing contributing to the FIE detection effect in conscious access. In Experiment 2, we tested how the recognizability of Mooney-like face stimuli influenced bCFS and rev-bCFS. Previous studies showed that attributing a specific meaning to a visual stimulus (compared to the same stimulus without such meaningful content and acquired perceptual structure) leads to an increase in perceptual dominance in a BR task (Yu & Blake, 1992). In a pre-post design, we tested detection effects for two-tone degraded face stimuli (i.e., Mooney faces; (Latinus & Taylor, 2005; Schwiedrzik et al., 2018)) in participants initially not aware of them being faces (pre-meaning reveal), but being informed about their meaning later (post-meaning reveal), as compared to participants not subjected to the pre-post reveal. We hypothesized the effect of stimulus recognition to involve conscious rather than unconscious processing, and thus slower disappearance timings in rev-bCFS for the group subjected to the pre-post reveal, but no effects in bCFS.

## 1. Materials and Methods

### 1.1 Experiment 1

#### 1.1.1 Participants

21 subjects were recruited for the study. They had normal or corrected-to-normal vision, no history of neurological diseases, and they were naïve concerning the research question. The sample size was estimated with a priori power analysis based on the FIE effect size *d* = .092 reported by Jiang and colleagues (2007). With such an effect size, for a one-tailed t-test with *alpha* = .05 and 99% power, the estimated sample size was 21 subjects. The study was approved by the Ethical Committee of the University of Turin (protocol n. 0486683), and participants gave informed consent to participate in the investigation.

#### 1.1.2 Apparatus, stimuli, and procedure

The experiments were programmed in Matlab (Release 2021b) using the Psychtoolbox (Brainard, 1997) functions and presented on a Q-BenX monitor (1920 x 1080 pixels resolution, 120Hz refresh rate). Participants sat in front of the screen at approximately 57 cm and in front of a chinrest with a built-in stereoscope that was adjusted for each participant to allow for stable binocular vision. The screen background was black, and two fusion squares (1.50° x 1.95°) with a black and white pixel contour (0.15°) were used for the binocular presentation of the stimuli. Stimuli (1.50° x 1.95°) consisted of 40 black and white faces (with neutral expression, half males and half females), matched in luminance and contrast, and cropped into oval shapes (Stein et al., 2017). High-contrast colorful masks were generated in Matlab. The order of the two tasks (bCFS and rev-bCFS) was counterbalanced across participants.

##### Breaking continuous flash suppression (bCFS) task

During the trial, one eye was exposed to a target face that was linearly ramped up in its contrast from 0 to 100% within the first 4s of the trial, while the other eye was exposed to a dynamic high-contrast mask flashing at 10 Hz of frequency. Mask contrast was 100% during the first 4 seconds of the trial, then its contrast linearly decreased from 100 to 0% in the remaining 8 seconds. Thus, each trial lasted for a maximum of 12 seconds, or until a response was made (*See Fig. 1*). Targets and masks covered the fusion squares, with the target being presented in the center. Participants were asked to maintain fixation to the central fixation cross, to avoid blinks during the trial, and to keep both eyes open during the experiment. Importantly, they were instructed to press the space bar as soon as they perceived any part of the target breaking suppression (i.e., when anything other than the mask became visible). They were instructed not to wait until they could identify the target image. Trials were separated by 2.1 s of inter-trial interval, and the task was composed of 160 randomized trials, with 80 trials containing upright face targets, and 80 trials containing their inverted counterparts (i.e., rotated by 180°). The target eye was also counterbalanced and randomized (in 80 trials the face was shown to the right eye and in the remaining 80 to the left eye).

**Figure 1:**
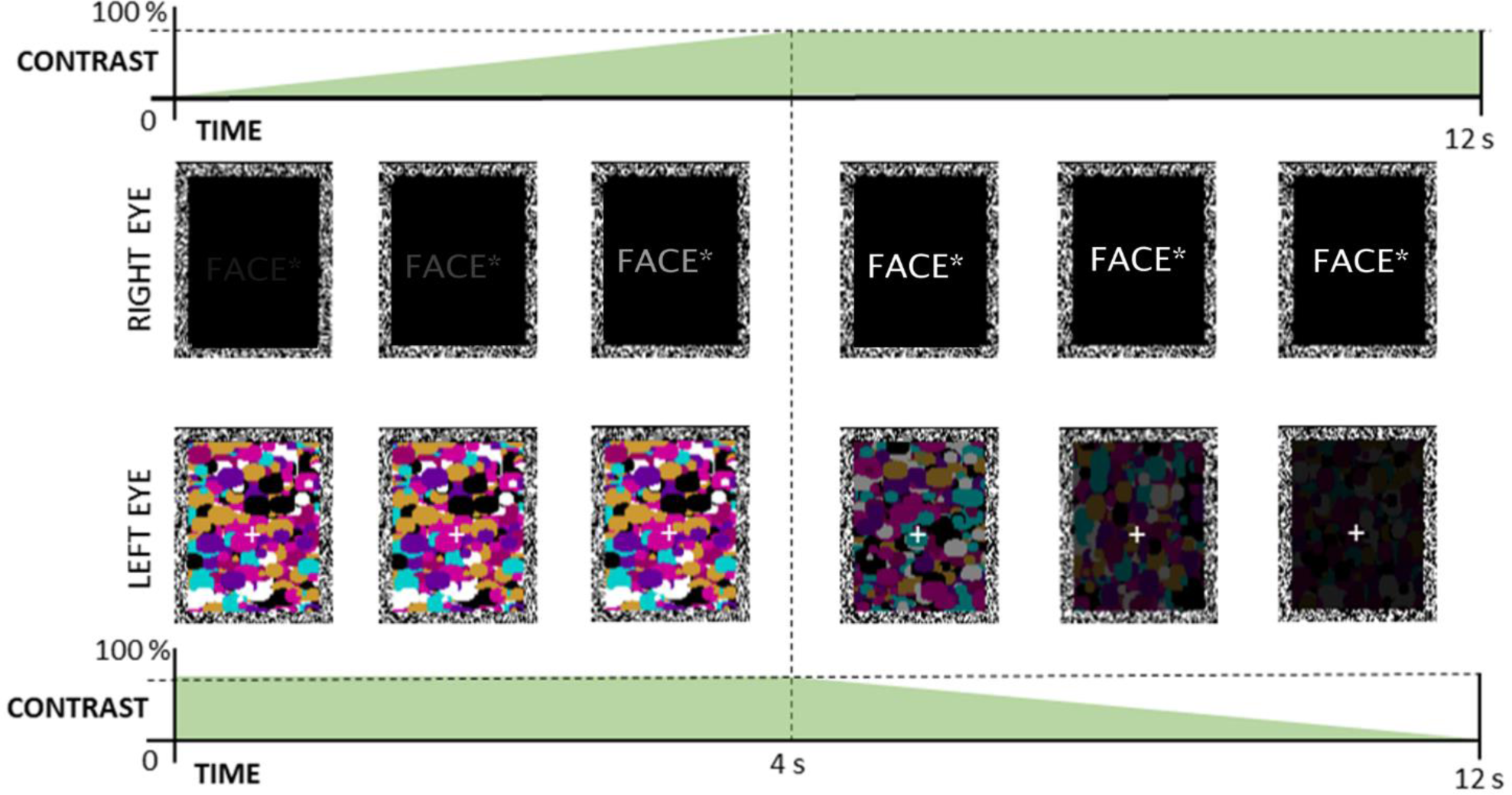
*Schematic representation of the bCFS trial*. After 2.1 seconds of inter-trial interval, a high-contrast mask (10Hz) was shown to one eye, and its contrast was decreased from 100 to 0% within 8 seconds after 4 seconds of trial. The target face was shown to the other eye, and its contrast was increased from 0 to 100% within the first 4s of the trial. Each trial lasted for a maximum of 12 seconds or until response (space bar) once participants detected the first cue of the target breaking the suppression. *Stimuli were real face images matched in contrast and luminance, as used in Stein et al. (2017).

After 12 trials of familiarization, the experiment began and lasted approximately 15 minutes, with a small break after 80 trials.

##### Reverse-breaking continuous flash suppression (rev-bCFS) task

The setup for the rev-bCFS experiment was almost identical to the bCFS experiment, with two main critical differences. First, contrast ramping phases were reversed: the target face started with 100% contrast for the first 4 s of the trial and decreased linearly from 100 to 0% over the next 8 s, while the masks started with 0% contrast and increased to 100% within the first 4s, remaining constant until response or the end of the trial (12 s; *see Fig. 2* for a schematic representation).

**Figure 2:**
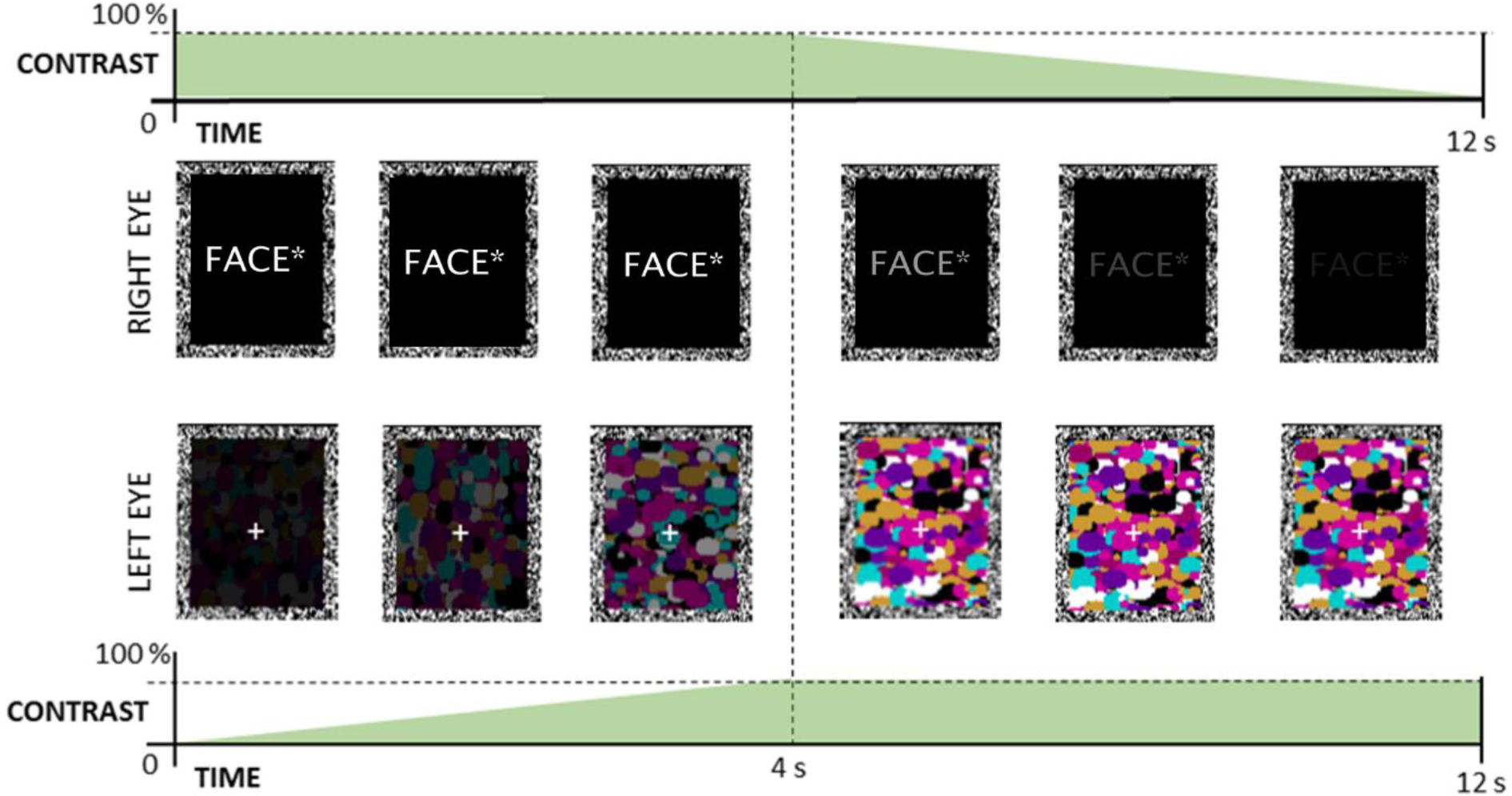
*Schematic representation of the rev-bCFS trial*. The target face, shown to one eye, was shown at full visibility in the first 4s of the trial, then its contrast was linearly decreased from 100 to 0% in the remaining 8s. On the other eye, the mask increased its contrast from 0 to 100% within the first 4s of the trial and remained stable until the end of the trial (12s) or until response. Participants had to press the space bar once the last cue of the target disappeared from their conscious percept. *Stimuli were real face images matched in contrast and luminance, as used in Stein et al. (2017).

Participants were asked to respond as quickly as possible to the disappearance of the target by pressing the space bar when the last part of the face became invisible.

#### 1.1.3 Statistical analysis

One participant reported unstable binocular perception and was excluded from the analysis. In the bCFS task, trials with response times lower than 300 ms (0.37% of the trials) were excluded (as this suggested that stimuli were not suppressed). To remove between-subjects variability that typically affects bCFS performance and is of no interest for the effect of the experimental manipulation, we used the latency-normalization procedure approach that has been used in other studies (Gayet et al., 2016; Gayet & Stein, 2017; Tsuchiya et al., 2006). This index has been shown to not only account for between-subject variability, but also to approximate RT differences to a normal distribution, similar to logarithmic transformations (that we also calculated), and to enhance RT differences caused by the experimental manipulation by reducing type II error rate. Parameters were calculated as in the work by Gayet and Stein (Gayet & Stein, 2017). The Latency-Normalized RT difference was thus scored as follows:

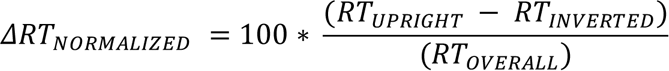

where *RT_UPRIGHT_* was defined as the median value of the upright condition, *RT_INVERTED_* as the median value of the inverted condition, *RT_OVERALL_* as the median RT’s average within each condition. Medians were used to account for the skewed distribution of the raw data. This index was calculated for the two tasks separately. Negative values indicate faster RT for upright faces.

Another normalized FIE index was calculated after log-transforming the raw RTs:

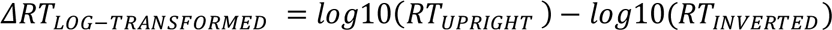

Planned t-test comparisons, analyzing differences between upright and inverted faces within each experiment, and comparing the face-inversion differences between the two experiments were conducted with JASP (JASP Team, 2016).

### 1.2 Results

*Breaking CFS task.* Median RTs were faster for upright (M = 3.09 s, SE = ± .27) than for inverted faces (M = 3.77, SE = ± .30), and this difference was statistically significant for both median RTs (*t*_(19)_ = -4.22, *p* < .001, Cohen’s *d* = .94) and log-transformed RTs (*t*_(19)_ = -4.32, *p* < .001, one-tailed, Cohen’s *d* = .96), indicating faster access to awareness for upright than for inverted faces. The negative latency-normalized index showed that upright face condition sped up RTs by 21.22% (SE = ± 4.8%). Thus, we replicated the standard FIE obtained in bCFS (raw difference = -678ms, SE = ± .16) with an effect size (Cohen’s *d* = .94) in line with other studies (Gayet & Stein, 2017; Jiang et al., 2007; Stein et al., 2016). *See Fig.3*.

**Figure 3:**
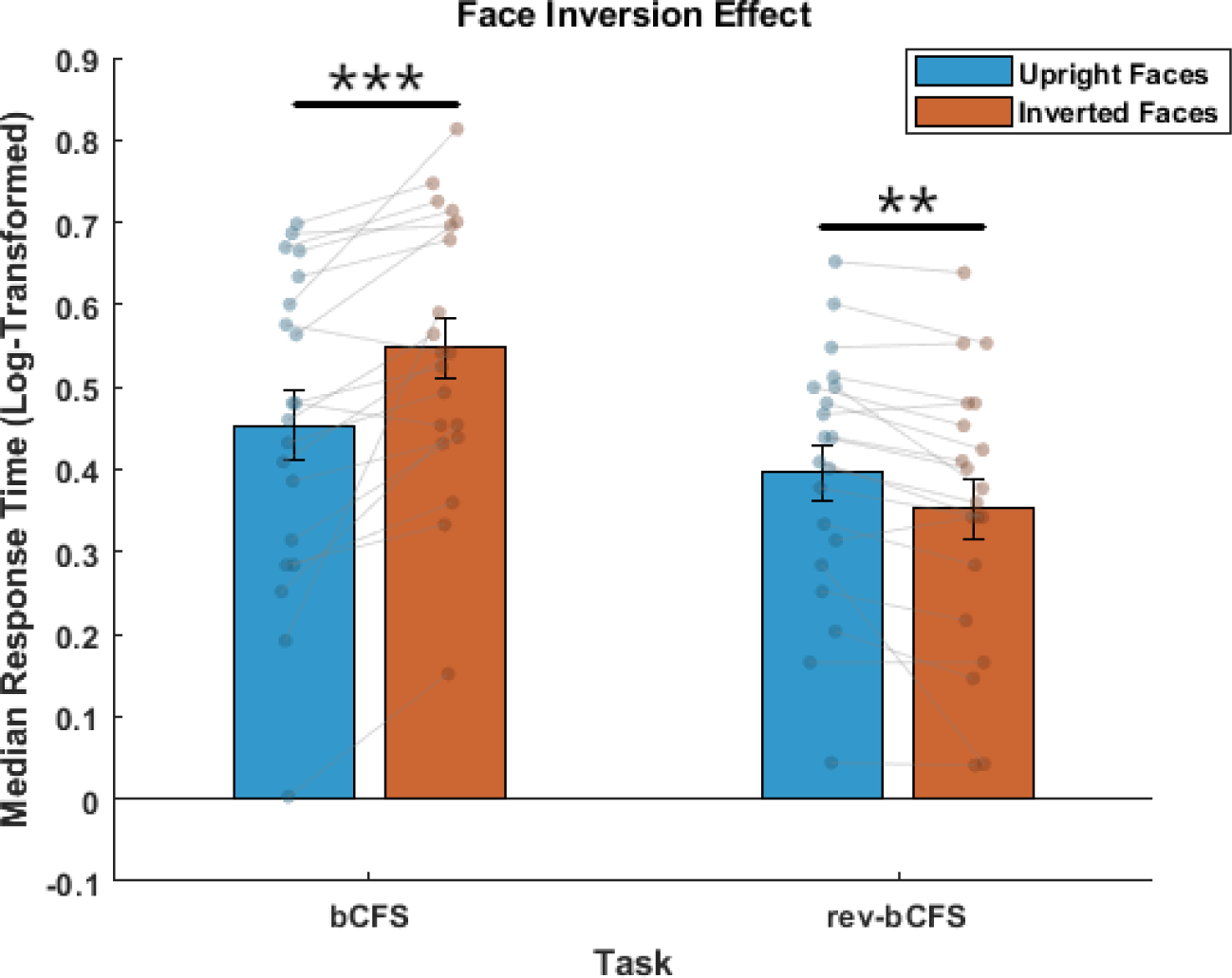
*Results of Experiment 1*. Median log-transformed RTs (and SE) from the breaking- continuous Flash Suppression task as a function of face orientation, reflecting the suppression time of the targets, and from the reverse-breaking Continuous Flash Suppression task, reflecting face disappearance/dominance. Points represent subject-based performance *\**p *< .05, \*\**p *< .01, \*\*\**p *< .001*.

*Reverse-breaking CFS task.* Median RTs for face disappearance from awareness were 2.52s (SE = ± .19), whereas upright median RTs were 2.63s (SE = ± .19), and 2.40s (SE = ± .06) for inverted exemplars. The face-inversion difference was 223ms (SE = ± .05). Data were normally distributed. Planned t-test comparison showed a significant difference between the two conditions in both raw (*t*_(19)_ = 4.35, *p* < .001, Cohen’s *d* = .97) and log-transformed (*t*_(19)_ = 3.40, *p* = .002, Cohen’s *d* = .76) data, with consistent effect sizes. These results indicate that upright faces, as compared to inverted ones, were slower in disappearing from awareness. The latency-normalization index was positive, showing that upright faces slowed down overall RTs by 9.9% (SE = ± 2.8%). *See Fig.3*.

*Tasks comparison.* We then compared the overall median RTs for faces within the two tasks. Overall RTs were longer in bCFS (*t*_(19)_ = 2.36, *p* = .029, Cohen’s *d* = .53), showing that suppression was faster than breaking suppression. Next, we compared the face inversion effect between the two tasks. As the face inversion effect in bCFS and rev-bCFS have opposite signs (i.e., faster RTs in bCFS and slower RTs in rev-bCFS, both indicating upright face prioritization), we reversed the bCFS FIE sign. The analysis revealed consistent results across the different metrics used, showing that the FIE was larger for bCFS compared to re-bCFS with raw median RTs (*t*_(19)_ = 2.51, *p* = .011, Cohen’s *d* = .56), log-transformed RTs (*t*(19) = 1.89, *p* = .037, Cohen’s *d* = .42), and latency- normalized effects (*t*(19) = 1.91, *p* = .036, Cohen’s *d* = .43). *See Fig.4*.

**Figure 4:**
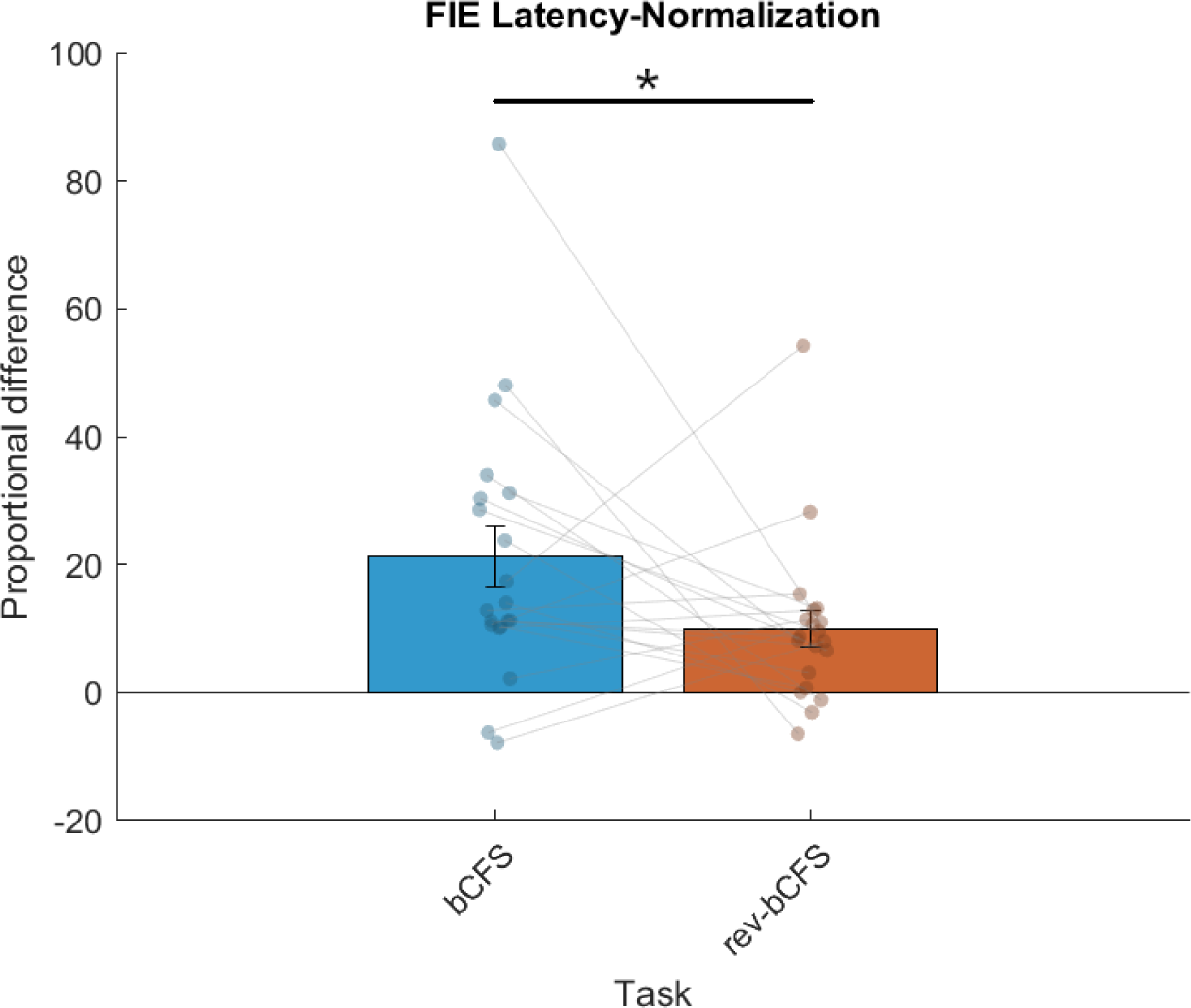
*Comparison of FIE indexes across the two tasks*. Latency-normalization indices for the face-inversion effect between the two tasks (with subjects’ performance and SE). *\**p *< .05, \*\**p *< .01, \*\*\**p *< .001*.

### 2 Experiment 2

#### 2.1.1 Participants

Fifty subjects with normal or corrected-to-normal vision and without previous history of neurological diseases were recruited for the study. The sample size was estimated with an a priori power analysis (through g*Power) based on a medium effect size (*f* = .25) for a rm Anova with between-within interactions for 2 groups, alpha = .05, and statistical power of 99%, resulting in 50 subjects. The study was approved by the Ethical Committee of the University of Turin (protocol n. 0486683), and Participants gave informed consent to participate in the study.

#### 2.1.2 Apparatus, stimuli, and procedure

Apparatus and task structure were identical to Experiment 1, except for the following changes. The two stimuli were an inverted exemplar of a two-tone “Mooney” face and a neutral and meaningless two-tone image (both 1.5° x 1.95°, see *Fig 5*). Verbal and written instructions for the two tasks (bCFS and rev-bCFS, as in Experiment 1) were provided to participants, who were then exposed to the two stimuli. The experimental group (N = 25) was told that they had to detect appearance and disappearance of the stimuli, and they performed one bCFS and one rev-bCFS blocks (counterbalanced) made of 36 trials each (18 each stimulus). After they completed the two blocks, they were told that one stimulus was an inverted face (pre/post meaning-reveal). The stimulus was then rotated by 180° (upright) until participants recognized the face, and rotated again, making sure that participants could still recognize the face while inverted. The same exposure time was dedicated to the neutral stimulus. After the meaning was revealed, they performed the two tasks again (counterbalanced). For the control group (N = 25), the procedure was the same, with the exception that the meaning reveals occurred before performing any block. As a control manipulation, we decided to expose participants to stimulus reveal, rather than not expose them to prevent spontaneous Mooney face recognition during the experiment.

**Figure 5:**
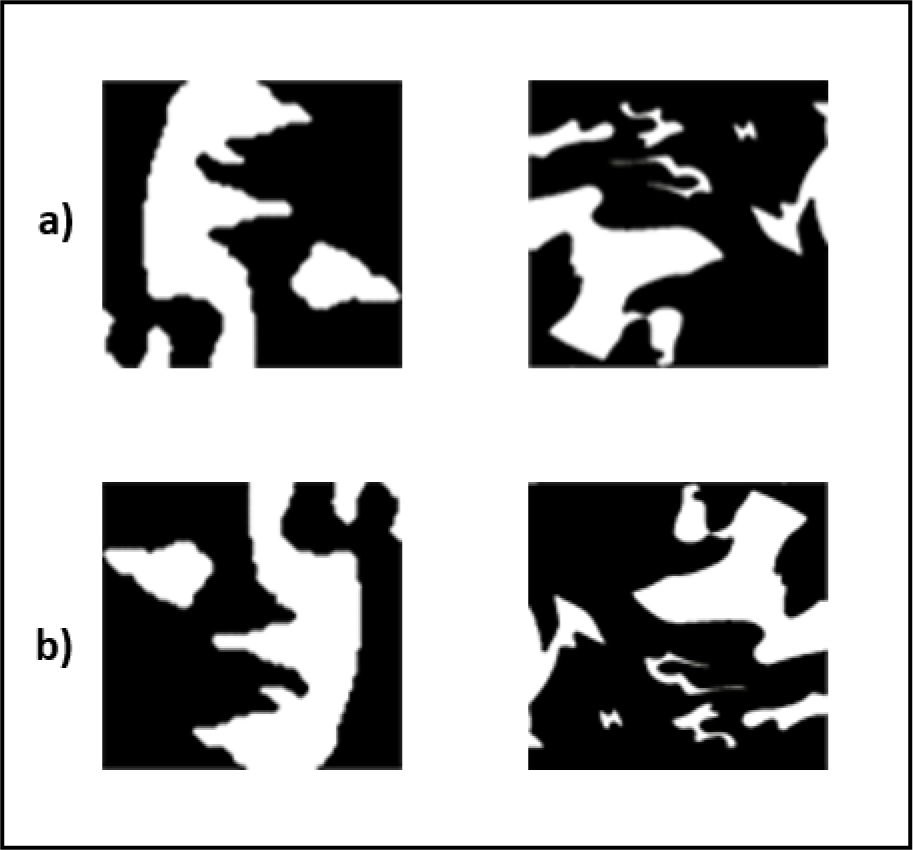
Targets of Experiment 2. a) the stimuli used in the two experiments, a two-tone inverted mooney face and a two-tone neutral stimulus. b) Shows the stimuli meaning reveal (same stimuli rotated by 180°), with participants recognizing the Mooney Face, which was rotated again for the post-experiment in the Experimental group, for the entire experiment in the control group. The same reveal time was dedicated to the neutral stimulus.

#### 2.1.3 Statistical analysis

In the bCFS task, trials with response time lower than 300 ms (0.32% of the trials) were excluded. We used the latency-normalization indices as in Experiment 1 by subtracting post blocks from pre blocks measures separately for each stimulus and task. The Latency-Normalized median differences were thus scored as follows:

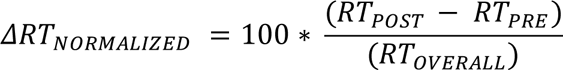

resulting in four measures (bCFS/rev-bCFS tasks, Mooney/Neutral stimuli). Negative scores indicate faster RT for stimuli presented in the post block. For task comparison, given that prioritization in bCFS and in rev-bCFS have opposite signs (i.e., faster RTs in bCFS and slower RTs in rev-bCFS), we flipped the sign for the bCFS latency normalized index.

### 2.2 Results

*Breaking CFS task.* A rm Anova was performed with the within-subject factor *Stimuli* (Mooney Face, Neutral) and the between-subject factor *Group* (Experimental, Control) on the latency normalized median differences. No significant effect was found, indicating that neither the stimulus nor the reveal manipulation or their interaction affected conscious access in post- vs. pre-reveal (*p* > .05). Equivalent results were found with median and log-transformed RTs.

*Reverse-breaking CFS task.* The same statistics were applied to latency normalized differences from the rev-bCFS task. Results revealed a main effect of *Group* (F_(1,48)_ = 9.29, *p* = .004, η_p_^2^ = .16), and t-test post-hoc comparisons showed that in the experimental group both stimuli (Mooney face and Neutral stimulus) persisted longer in visual awareness in post- vs. pre-reveal (*t*_(48)_ = -3.05, *p* = .004, Cohen’s *d* = .77). No other significant effects were found (*p* > .05). Similar results were confirmed with log-transformed (*t*_(48)_ = -3.04, *p* = .004, Cohen’s *d* = .78) and raw differences (*t*_(48)_ = -2.35, *p* = .023, Cohen’s *d* = .58).

#### Tasks comparison

The performance of the two tasks was directly compared with a rm Anova with the within factors *Stimuli* (Mooney Face, Neutral), *Task* (bCFS, rev-bCFS) and the between factor *Group* (Experimental, Control) for latency normalized differences. Results showed a significant interaction between *Group*\**Task* (F_(1,48)_ = 6.14, *p* = .017, η_p_^2^ = .11; see Fig.6). No other significant difference was found. Post-hoc t-test comparisons revealed that only the Experimental group showed longer persistence of both stimuli in the rev-bCFS only (*t*_(48)_ = -3.05, *p* = .004, Cohen’s *d* = .77).

**Figure 6:**
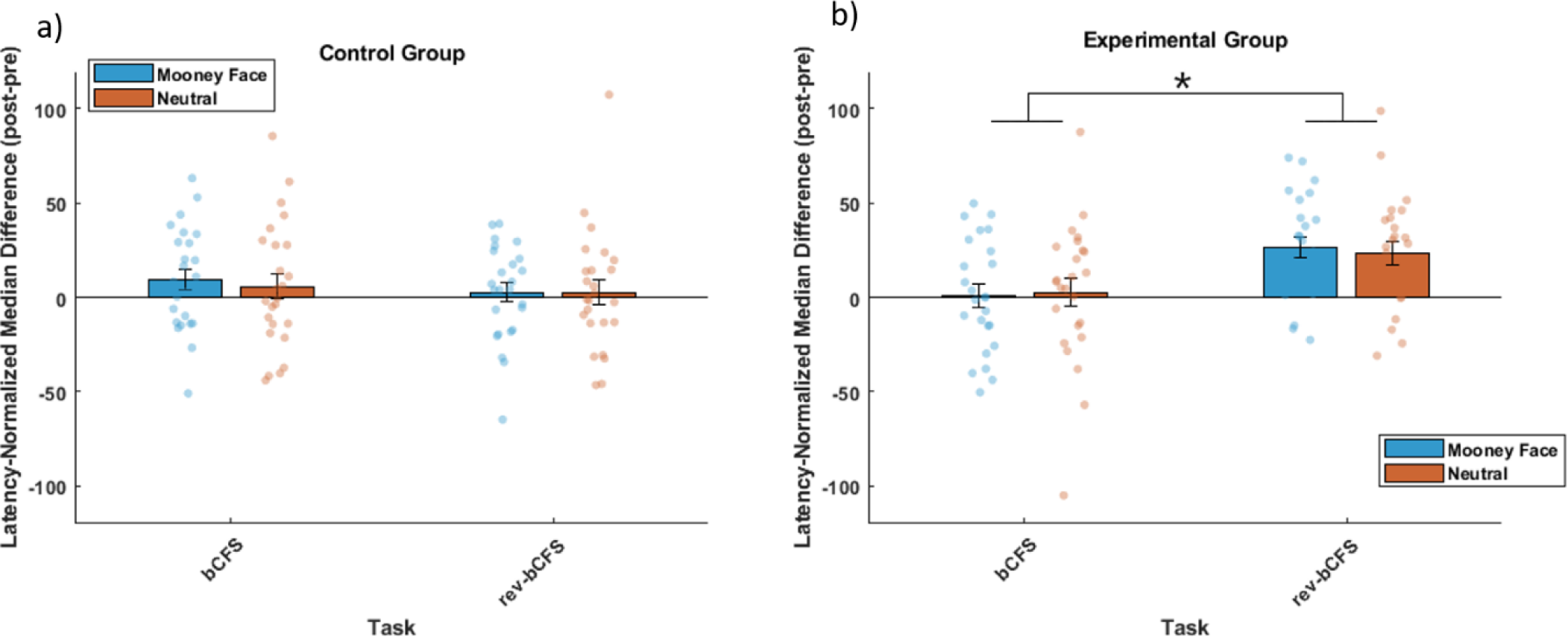
Results Experiment 2: Results of Experiment 2 showing the Latency-Normalized indices (post-pre) in control (a) and experimental (b) participants, with subjective points and SE. \**p* < .05, \*\**p* < .01, \*\*\**p* < .001.

## 3 Discussion

We introduced reverse-breaking continuous flash suppression (rev-bCFS) as a novel approach to measure and control for conscious influences on suppression times recorded with the popular breaking continuous flash suppression (bCFS) paradigm. In two experiments using face stimuli we sought to disentangle conscious and unconscious influences on detection effects by comparing conscious access (bCFS) and disappearance (rev-bCFS) using face stimuli. Specifically, we tested the face inversion effect (FIE; Experiment 1) and the effect of stimulus recognizability using two-tone Mooney-like stimuli (Experiment 2). Based on the assumption that rev-bCFS captures conscious effects, we reasoned that detection effects that are larger in standard bCFS than in rev- bCFS would reflect an additional contribution of unconscious processing occurring during suppression. Indeed, in Experiment 1 we found that the FIE was greater in bCFS than in rev-bCFS, suggesting that unconscious processing contributed to the advantage of upright over inverted faces in accessing awareness. In Experiment 2, we found that assigning a meaning to two-tone stimuli slowed disappearance timings in rev-bCFS but had no effect on bCFS. This effect was not specific to two-tone Mooney-like faces, but similarly present for a meaningless neutral two-tone stimulus, indicating that the effect was driven by top-down attention affecting conscious but not unconscious processing.

The greater FIE in bCFS suggests that there is a distinct contribution of unconscious processing to the effect that goes above and beyond the conscious prioritization of upright faces seen in rev-bCFS. The putative neural mechanisms underlying such processing are still partially unclear; whereas some findings highlight the role of a subcortical pathway for fast and coarse face processing (Stein, Peelen, et al., 2011; G. Zhou et al., 2010), others highlighted the importance of extrastriate cortical activity in the FFA for the FIE (Yovel & Kanwisher, 2005) that can survive interocular suppression (Jiang & He, 2006). Such preferential processing for upright faces has been hypothesized to reflect an innate perceptual predisposition for faces, being present at early stages of the lifespan (Farroni et al., 2005; McKone et al., 2007), as well as effects of visual familiarity acquired during the lifespan (Laguesse et al., 2012; Stein et al., 2014).

The results of Experiment 2 were quite different: an effect of revealing stimulus meaning was observed only in rev-bCFS, in which disappearance timings were slower in the group that was informed about the meaning of the Mooney-like face stimulus. The effect was not specific to the two-tone face stimulus but similarly seen for the meaningless two-tone neutral stimulus. This suggests that this effect was driven by top-down attention, with participants deploying greater attention to the previously meaningless stimuli after they had been informed about their meaning. Indeed, attention has been shown to influence dominant phases of stimuli in BR (Chong & Blake, 2006; Mitchell et al., 2004; Paffen & Alais, 2011; van Ee et al., 2005). Crucially, this effect was not observed in bCFS, suggesting that it involved conscious mechanisms of top-down attention but no unconscious processing occurring under suppression.

We suggest that rev-bCFS could represent a useful approach for the study of timings for visual awareness by measuring and controlling for conscious influences on detection effects, and an improvement over the standard non-CFS control conditions, given that bCFS and rev-bCFS share interocular suppression, similar perceptual dynamics, uncertainty, and require a similar detection task around the subjective threshold to awareness. Recently and independently from the present work, a similar approach has been used, measuring appearance from suppression and disappearance from dominance for different images categories (e.g., faces vs. objects) during CFS in a continuous cycle (Alais et al., 2024). These authors found that differences between images categories (e.g., faces requiring lower contrast to appear and to disappear than objects) were similar for appearance and disappearance. In line with our logic here, this was interpreted as showing that high-level stimulus properties such as category membership do not influence suppression. These findings appear inconsistent with standard hybrid models of binocular rivalry that postulate high-level stimulus effects on suppression (Hesse & Tsao, 2020; Tong et al., 2006), as well as with our observation of a larger FIE during bCFS than rev-bCFS, which suggests an effect of configural stimulus properties on suppression. One reason for this apparent discrepancy is that although Alais and colleagues (2024) carefully matched certain low-level properties of their stimuli, it is difficult to interpret detection of physically different target stimuli during CFS (Stein, 2019). Here, we therefore compared physically identical stimuli, changing only their spatial orientation (Experiment 1) or their ascribed meaning (Experiment 2). In addition, the continuous presentation cycle employed in this previous study may have resulted in adaptation effects that are prevented by comparing bCFS and rev-bCFS using separate trials with fixed presentation durations.

Before concluding, some cautionary notes are in order: we are not suggesting that the rev- bCFS paradigm can provide unequivocal evidence that unconscious processing differences contributed to detection effects. The present approach relies on the assumption that conscious and unconscious processes determine detection speed, and that rev-bCFS captures all conscious processes that could drive bCFS detection effects. If this assumption is not met, larger effects in bCFS may be driven by conscious factors other than those captured by rev-bCFS. Although we cannot exclude this possibility here, we believe that the rev-bCFS represents a promising step towards disentangling conscious and unconscious effects on visual awareness timings. To unequivocally establish that a detection effect was caused by unconscious factors, accuracy-based dissociation paradigms are required. For example, using backward masking, recent work (Stein & Peelen, 2021) measured detection differences (e.g., better detection of upright than inverted faces) while simultaneously ensuring that subjects had no access to the key manipulation driving these detection effects (e.g., they could not distinguish between upright and inverted faces). However, such accuracy-based dissociation approaches are challenging to combine with CFS, given its inter- and intraindividual variability, and its unpredictability in strength and depth of suppression.

Moreover, objective awareness measures such as those employed by Stein and Peelen (2021) may be too conservative and thus risk underestimating the extent of unconscious processing. The rev- bCFS paradigm, instead, relies on intuitively appealing subjective measures of awareness that aim to capture the subjective appearance and disappearance of contents in visual consciousness.

Finally, another advantage of rev-bCFS is that it can be used to measure conscious effects on the dynamics of interocular suppression that are not influenced by suppression times as in the classic BR alternation cycle. Indeed, in BR it cannot be estimated whether longer dominance times reflect prioritization during conscious perception or prioritization outside of awareness, with stimuli spending less time in the suppression phases. Thus, with BR it is difficult to disentangle the contribution of conscious and unconscious processing on visual awareness. Furthermore, dominance and suppression phases are known to rely on distinct brain mechanisms that can differentially affect interocular suppression timings (Blake & Logothetis, 2002), and BR and CFS are characterized by differential depth of suppression (Tsuchiya et al., 2006). For that reason, the combined use of bCFS and rev-bCFS could represent an improvement over BR to investigate dominance and suppression timings independently, as well as a useful tool to study differential brain dynamics for an appearing or disappearing stimulus from awareness that might entail implications for current theories on the neural correlates of consciousness (Seth & Bayne, 2022).

In conclusion, we introduced rev-bCFS as a practical and straightforward approach to control for conscious effects on detection measured with bCFS. The combination of bCFS and rev- bCFS may represent a promising experimental paradigm for disentangling conscious and unconscious influences on access to awareness and thus better isolate unconscious processing than standard non-CFS control conditions. This approach may prove fruitful in future investigations testing the scope and limits of unconscious processing under interocular suppression.

## Acknowledgments

We thanks Trombetti L. for help with data collection.

## Authors Contributions

T.C. and T.S. conceived the original idea; T.C. collected and analyzed the data; T.C., L.P. and T.S. designed the study, interpreted the data and drafted the paper.

